# Vaccination with single plasmid DNA encoding IL-12 and antigens of severe fever with thrombocytopenia syndrome virus elicits complete protection in IFNAR knockout mice

**DOI:** 10.1101/787879

**Authors:** Jun-Gu Kang, Kyeongseok Jeon, Hooncheol Choi, Yuri Kim, Hong-Il Kim, Hyo-Jin Ro, Yong Bok Seo, Jua Shin, Junho Chung, Yoon Kyung Jeon, Yang Soo Kim, Keun Hwa Lee, Nam-Hyuk Cho

**Affiliations:** Department of Microbiology and Immunology, Seoul National University College of Medicine, Seoul, Republic of Korea; Department of Biomedical Sciences, Seoul National University College of Medicine, Seoul, Republic of Korea; SL-VAXiGEN Inc., Seongnam, Gyeonggi, Republic of Korea; Department of Biochemistry and Molecular Biology, Seoul National University College of Medicine, Seoul, Republic of Korea; Department of Pathology, Seoul National University College of Medicine, Seoul, Republic of Korea; Department of Infectious Diseases, Asan Medical Center, University of Ulsan College of Medicine, Seoul, Republic of Korea; Department of Microbiology and Immunology, Jeju National University School of Medicine, Jeju, Republic of Korea; Institute of Endemic Disease, Seoul National University Medical Research Center and Bundang Hospital, Seoul, Republic of Korea

**Keywords:** Severe fever with thrombocytopenia syndrome, DNA vaccine, T cells, protective immunity

## Abstract

Severe fever with thrombocytopenia syndrome (SFTS) is an emerging tick-borne disease caused by SFTS virus (SFTSV) infection. Despite a gradual increase of SFTS cases and high mortality in endemic regions, no specific viral therapy nor vaccine is available. Here, we developed a single recombinant plasmid DNA encoding SFTSV genes, Gn and Gc together with NP-NS fusion antigen, as a vaccine candidate. The viral antigens were fused with Fms-like tyrosine kinase-3 ligand (Flt3L) and IL-12 gene was incorporated into the plasmid to enhance cell-mediated immunity. Vaccination with the DNA provides complete protection of IFNAR KO mice upon lethal SFTSV challenge, whereas immunization with a plasmid without IL-12 gene resulted in partial protection. Since we failed to detect antibodies against surface glycoproteins, Gn and Gc, in the immunized mice, antigen-specific cellular immunity, as confirmed by enhanced antigen-specific T cell responses, might play major role in protection. Finally, we evaluated the degree of protective immunity provided by protein immunization of the individual glycoprotein, Gn or Gc. Although both protein antigens induced a significant level of neutralizing activity against SFTSV, Gn vaccination resulted in relatively higher neutralizing activity and better protection than Gc vaccination. However, both antigens failed to provide complete protection. Given that DNA vaccines have failed to induce sufficient immunogenicity in human trials when compared to protein vaccines, optimal combinations of DNA and protein elements, proper selection of target antigens, and incorporation of efficient adjuvant, need to be further investigated for SFTSV vaccine development.

**Author summary:** Severe fever with thrombocytopenia syndrome (SFTS) is an emerging tick-borne infection endemic to East Asia including China, Korea, and Japan. Gradual rise of disease incidence and relatively high mortality have become a serious public health problem in the endemic countries. In this study, we developed a recombinant plasmid DNA encoding four antigens, Gn, Gc, NP, and NS, of SFTS virus (SFTSV) as a vaccine candidate. In order to enhance cell-mediated immunity, the viral antigens were fused with Flt3L and IL-2 gene was incorporated into the plasmid. Immunization with the DNA vaccine provides complete protection against lethal SFTSV infection in IFNAR KO mice. Antigen-specific T cell responses might play a major role in the protection since we observed enhanced T cell responses specific to the viral antigens but failed to detect neutralizing antibody in the immunized mice. When we immunized with either viral glycoprotein, Gn protein induced relatively higher neutralizing activity and better protection against SFTSV infection than Gc antigen, but neither generated complete protection. Therefore, an optimal combination of DNA and protein elements, as well as proper selection of target antigens, might be required to produce an effective SFTSV vaccine.

## Introduction

Severe fever with thrombocytopenia syndrome (SFTS) is an emerging tick-borne infectious disease caused by SFTS virus (SFTSV), belonging to the *Phenuiviridae* family of *Bunyavirales* [1, 2]. The genome of SFTSV is composed of three segmented RNAs: large (L) segment encoding RNA-dependent RNA polymerase (RdRp), medium (M) encoding the envelop glycoproteins, Gn/Gc, and small (S) encoding the nucleocapsid and nonstructural proteins (NP and NS) [1]. Clinical manifestations include fever, gastrointestinal symptoms, leukocytopenia, and thrombocytopenia [3, 4]. Disease mortality of SFTS patients has been estimated to be 5 ∼ 20% [3]. Even though the majority of SFTS cases has been reported from China [3], Korea [4], and Japan [5], SFTSV infections in southern Asia, including Vietnam, have been recently reported in a retrospective survey [6]. Currently, no specific viral therapy nor vaccine is available. An effective vaccine is needed to combat its relatively high mortality, especially in elderly patients, and spread of SFTSV between humans [7, 8].

Vaccine development for SFTS is at an early discovery phase and there have only been a few studies on vaccine candidates using animal infection models [8-11]. Immunization of NS antigen with Freund’s adjuvant in C57BL/6 mice, which are naturally resistant to SFTSV but partially mimic human infections [12], failed to enhance viral clearance, although it induced high titer of anti-NS antibodies and significantly elevated IFN-γ levels in sera upon viral challenge [9]. Vaccination of plasmid DNAs encoding NP and NS peptides also enhanced antigen-specific cellular immunity of T cells, such as IFN-γ and TNF-α secreting CD4^+^ and CD8^+^ T cells, in BALB/c mice when applied by nano-patterned microneedles [10]. However, the protective effect of cellular immunity induced by DNA vaccination was not confirmed by *in vivo* infection with SFTSV. Recently, Dong F. *et al*. reported that a single dose of live attenuated recombinant vesicular stomatitis virus (rVSV) vaccine expressing the SFTSV Gn/Gc glycoproteins elicited high titers of protective neutralizing antibodies in both wild type and interferon α/β receptor knockout (IFNAR KO) mice [11]. They clearly showed that a single dose rVSV carrying SFTSV Gn/Gc could provide complete protection against lethal challenge with SFTSV in young and old IFNAR KO mice, a promising infection model for severe human infection of SFTSV [11, 13, 14]. Nevertheless, the potential role of cellular immunity against the viral antigens in complete protection was not examined in this study, although adoptive transfer of immune sera from mice immunized with rVSV-Gn/Gc to naïve IFNAR KO mice provide partial (∼ 60%) protection against lethal SFTSV challenge [11].

In this study, we developed a recombinant plasmid DNA (pSFTSV) encoding extracellular domains of Gn and Gc, and NP-NS fusion antigen as a DNA vaccine candidate. We examined whether it could provide protective immunity against lethal SFTSV infection in IFNAR KO mice. In order to facilitate the processing and presentation of the SFTSV antigens by dendritic cells (DCs) and enhance antigen-specific T cell responses, these recombinant antigens were fused with Fms-like tyrosine kinase-3 ligand (Flt3L) [15, 16]. Moreover, we generated a recombinant DNA encoding IL-12α and β in addition to the recombinant viral antigens (pSFTSV-IL12) to further enhance cell-mediated immunity [17]. Vaccination of pSFTSV-IL12 provided complete protection of IFNAR KO mice upon lethal SFTSV challenge, whereas immunization with pSFTSV elicits only partial protection, indicating that antigen-specific cellular immune responses enhanced by co-expression of IL-12 could play a significant role in protection against lethal SFTSV infection. We confirmed significantly higher levels of Gn and NP-specific CD4^+^ and CD8^+^ T cell responses in mice vaccinated with pSFTSV-IL12 when compared to those in mock vector-immunized mice. Therefore, our results indicate that enhanced antigen-specific T cell immunity against multiple SFTVS antigens by DNA vaccination could be a promising direction for developing an effective vaccine against SFTSV infection.

## Methods

### Ethics statement

Animal experiments were conducted in an Animal Biosafety Level 3 facility at Seoul National University Hospital and International Vaccine Institute. This study was approved by the Seoul National University Hospital and International Vaccine Institute Institutional Animal Care and Use Committee (SNUH IACUC No. 15-0095-C1A0 and IVI IACUC No. 2018-018) and conducted in strict accordance with the recommendations in the National Guideline for the care and use of laboratory animals.

### Preparation of SFTSV DNA plasmid

In order to generate plasmids for DNA vaccination, genes encoding ectodomains of Gn, Gc (from Genebank accession no. AJO16082.1), and NP/NS fusion protein (from Genebank accession no. AJO16088.1 and AKI34298.1, respectively) were synthesized (GenScript, Piscataway, NJ, USA) and cloned into pGX27 vector (Genexine, Seongnam, Republic of Korea). All these genes were fused with signal peptide of tissue Plasminogen Activator (tPA, Uniport no. P00750) and Flt3L (Uniport no. P49771) in their N-terminus [15] (pSFTSV, Fig 1). In addition, murine IL-12α and β genes (Uniport no. P43432) [17] were also synthesized (GenScript) and cloned into pSFTSV (pSFTSV-IL-12, Fig 1).

**Fig 1.**
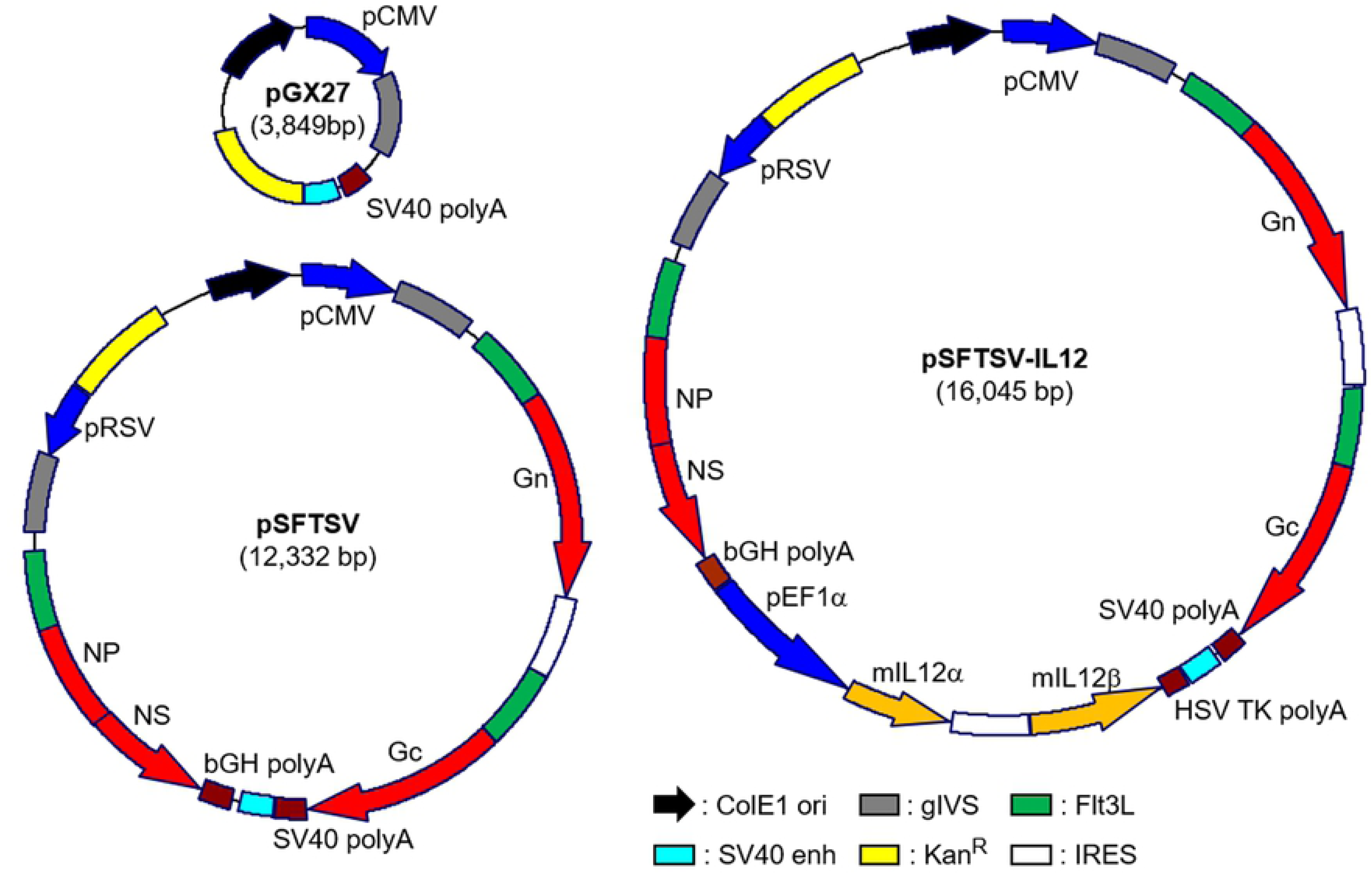
Schematic diagrams of SFTSV DNA vaccines used in this study. pSFTSV was constructed by inserting genes encoding ectodomains of Gn and Gc, and NP-NS fusion protein into the pGX2 vector. The viral antigens are preceded by the secretory signal sequence of tPA and the extracellular domain of Flt3L. pSFTSV-IL-12 includes additional genes encoding murine IL-12α and β to promote cellular immunity. ColE1, ColE1-type bacterial origin of replication; gIVS, rabbit β-globin intervening sequence; Kan^R^, kanamycin resistance gene; IRES, internal ribosome entry site.

### Gene expression of plasmid DNAs

To confirm the expression of cloned genes in pSFTSV and pSFTSV-IL12, HEK 293T cells (ATCC CRL-1573, Manassas, VA, USA) were transfected with pGX27, pSFTSV, or pSFTSV-IL12 plasmid using the polyethylenimine (PEI) transfection method [18]. Briefly, plasmids and PEI were diluted with Opti-MEM media (Gibco, Gaithersburg, MD, USA) and incubated with HEK 293T cells. After 4 h of incubation, media were changed with Dulbecco’s Modified Eagle medium (DMEM) (Gibco) containing 10% fetal bovine serum (FBS) and incubated in humidified CO_2_ atmosphere at 37°C. After 48 h of culture, cells and culture supernatants were harvested. Cells were washed with phosphate-buffered saline (PBS) and lysed with NP-40 lysis buffer (1% NP-40 in 50 mM Tris-HCl, pH 8.0, 150 mM NaCl) containing protease inhibitor cocktail (Sigma-Aldrich, St. Louis, MO, USA). Lysates and supernatants were stored at -80°C until use.

For quantification of Flt3L-fused proteins and mouse IL-12 in the supernatants of transfected HEK293T cells, human Flt3L and mouse IL-12p70 Quantikine ELISA kits (R&D systems, Minneapolis, MN, USA) were used according to manufacturer’s instructions. Expression of Gc, Gn, and NP/NS fusion protein in the culture supernatants and cell lysates were examined by immunoblotting using rabbit anti-Gn, Gc (NBP2-41153 and NBP2-41156, NOVUS Biologicals, Centennial, Colorado, USA), and NP (produced by custom polyclonal antibody production service via Abclon, Seoul, Republic of Korea) antibodies, respectively. Each sample was separated on 8% polyacrylamide gels and transferred onto 0.45 µm PVDF membranes (Merck, Darmstadt, Germany). PVDF membranes were blocked with PBS containing 0.05% Tween 20 (PBST) and 5% skim milk at room temperature for 2 h. The membranes were sequentially incubated with primary and secondary antibodies conjugated with horse radish peroxidase (HRP, Invitrogen, Waltham, MA, USA). The membranes were then visualized using a LAS-4000 system (Fujifilm, Tokyo, Japan).

### Preparation of recombinant proteins

The gene encoding the NP of the KASJH strain (Genbank Accession No. KP663733) was cloned into the pET28a (+) vector (Novagen, Gibbstown, NJ, USA). NP protein was then purified from *E. coli* strain BL21 (DE3) harboring the recombinant plasmid. Following induction with 0.1 mM isopropyl β-D-thiogalactoside (IPTG) for 18 h at 16°C, the protein was purified using HisTrap HP histidine-tagged protein columns (GE healthcare, Chicago, IL, USA) according to the manufacturer’s instruction. Recombinant Gn and Gc glycoproteins fused to Fc region of human immunoglobulin heavy chain were purified as previously described [19]. Briefly, vectors cloned with gene encoding Gn or Gc were transfected into HEK293F cells (Thermo Fisher Scientific, Waltham, MA, USA) using polyethylenimine. The transfected cells were cultured in FreeStyle™ 293 expression medium (Gibco) for 6 days. Overexpressed recombinant proteins in supernatants were purified by AKTA start affinity chromatography system HiTrap Mabselect (GE Healthcare) according to the manufacturer’s instructions (S1 Fig).

### Enzyme-linked immunosorbent assays (ELISA)

To determine the antibody titers specific to Gc, Gn, and NP in sera of immunized mice, immunoassay plates (96-well plates; Nunc, Rochester, NY, USA) were coated with 100 ng/well of purified antigens at 4°C overnight. His-tagged Gn and Gc recombinant proteins were purchased from Immune Technology Co. (New York, NY, USA). His-tagged NP recombinant protein was purified as mentioned above. After antigen coating, the immunoassay plates were blocked for 2 h at room temperature with PBST containing 5% skim milk. 100 μl of serially-diluted serum samples were incubated for 1 h at room temperature and subsequently detected using HRP-conjugated goat anti-mouse IgG (Santa Cruz Biotechnology, Santa Cruz, CA, USA). 3,3’,5,5’-tetramethylbenzidine peroxidase substrate solution (KPL, Gaithersburg, MD, USA) was then added to develop color for 7 min, and the reaction was stopped by the addition of 1 M H_3_PO_4_ solution. Absorbance was measured at 450 nm using a microplate reader (TECAN, Mannedorf, Switzerland).

### Flow cytometric analysis

Spleen cells were released into RPMI 1640 media (Gibco) by mincing the spleen through a 70 μm cell strainer (BD Biosciences, San Jose, CA, USA). After lysis of red blood cell with a Red Blood Cell Lysing Buffer Hybri-Max™ (Sigma, St. Louis, MO, USA), splenocytes were cultured for 18 h in RPMI media containing 10% FBS (Gibco) and 1% penicillin/streptomycin (Gibco) in the presence of 10 μg of purified Gn or NP antigens in 96 well V-bottomed culture plates (Nunc, Roskilde, Denmark). For intracellular detection of IFN-γ, splenocytes (2 × 10^6^ cells/well) were treated with 1 μg of Golgiplug (BD Bioscience) for the final 6 h. Cells were then blocked with ultra-block solution (10% rat sera, 10% hamster sera, 10% mouse sera (Sigma), and 10 μg/ml of anti-CD16/32 (2.4G2) (BD Pharmingen, Franklin Lakes, NJ, USA), followed by staining with anti-CD3 (145-2c11) (BD Biosciences), CD4 (RM4-59) (BD Biosciences) and CD8 (53-6.7) (Biolegend, San Diego, CA, USA) antibodies conjugated to different fluorescent dyes. After surface staining, splenocytes were fixed and permeabilized with Cytofix/Cytoperm kit (BD Bioscience) and incubated with anti-IFN-γ antibody (XMG1.2) (BD Pharmingen). The stained cells were analyzed on a CytoFLEX S flow cytometer (Berkman Coulter Inc, Brea, CA, USA). Flow cytometry data were analyzed using FlowJo software version 10.6 (Tree Star, Ashland, OR, USA). Gating strategies for the flow cytometric analyses are summarized in S2 Fig.

### Preparation of SFTSV

SFTSV (Genbank accession no. MN329148-MN329150) isolated from a Korean SFTS patient was propagated in Vero E6 cells (ATCC CRL-1586). The supernatant of infected cells was harvested at 5 d after infection and stored at - 80°C after filtering with 0.45 μm syringe. Focus-forming unit (FFU) of SFTSV was determined by plaque assay using methylcellulose media [20]. Briefly, the filtered supernatants were serially diluted and added to a monolayer of Vero E6 cells and incubated for 1 h at 37°C. Viral supernatants were removed and cells were incubated under an overlay media (DMEM supplemented with 5% FBS and 1% methylcellulose) at 37°C for 7 days. Cells were fixed with 4% paraformaldehyde (Intron, Seongnam, Republic of Korea) and 100% methanol (Merck, Darmstadt, Germany). The SFTSV foci were detected using rabbit anti-SFTS NP antibody (Abclon) and goat anti-rabbit IgG secondary antibody conjugated with alkaline phosphatase (Thermo Fisher Scientific). Viral plaques were visualized by incubation with NBT/BCIP solution (Roche, Mannheim, Germany).

### Neutralizing antibody assay

To evaluate the neutralizing activity of immune sera, a focus reduction neutralization titer (FRNT) assay was performed using immunized mice sera [21]. SFTSV (0.0001 multiplicity of infection) was pre-incubated with serially diluted sera from mice at 4°C for 1 h. The mixture of virus and sera was added onto a monolayer of Vero E6 cells in 24-well plate. After incubation for 2 h, supernatant of cells were removed and cells were cultured under an overlay media at 37°C for 7 days. Viral foci were visualized as described above. The percentage of focus reduction was calculated as [(No. of plaques without antibody) – (No. of plaques with antibody)] / (No. of plaques without antibody) x 100. 50% focus reduction neutralization titers (FRNT_50_) were calculated by a nonlinear regression analysis (log[inhibitor] versus normalized response method) embedded in GraphPad Prism Software v5.01 (GraphPad Software; https://www.graphpad.com).

### Immunization of mice and SFTSV challenge

Six to eight-week-old female Interferon α/β receptor knockout (IFNAR KO, B57BL/6) mice [22] were used for immunization and challenge tests. They were housed and maintained in the specific pathogen-free facility at Seoul National University College of Medicine. Mice were intramuscularly immunized by electroporation using Orbijector EP-I model (SL Vaxigen Inc., Seongnam, Republic Korea) in the hind leg three times at two-week intervals. 4 μg of purified pGX27, pSFTSV, or pSFTSV-IL12 in 100 μl of PBS was used for each immunization. Mice were also subcutaneously immunized with 20 μg of Gn-Fc or Gc-Fc protein absorbed in aluminum hydroxychloride (Alhydrogel^®^ adjuvant 2%, InvivoGen, Hong Kong). Mice sera were collected from immunized mice at one week after the third immunization to determine the levels of specific antibody titers. Two weeks after the final immunization, mice were subcutaneously challenged with 1 × 10^5^ FFU of SFTSV. Body weight and mice survival were monitored until surviving mice fully recovered. Blood and tissues of mice were collected at the indicated time after viral challenge and applied for viral quantitation using qRT-PCR, or hematological analysis.

### Quantitative reverse transcript – polymerase chain reaction (qRT-PCR)

Total RNA was extracted from the plasma of SFTSV-infected mice using Trizol LS Reagent (Life Technologies, Carlsbad, CA, USA) according to the manufacturer’s instruction. Total RNA was reverse transcribed into cDNA using HiSenScript™ RH (-) RT Premix kit (Intron, Seongnam, Republic of Korea). cDNA was quantified using TaqMan Universal Master Mix 2 (Applied Biosystems, Waltham, MA, USA). qRT-PCR was performed on BioRad CFX connect real-time system (Bio-Rad, Hercules, CA, USA) under following conditions: uracil-N-glycosylase incubation at 50°C for 2 min, polymerase activation at 95°C for 10 min, denaturation at 95°C for 15 s, annealing and extension at 53°C for 1 min. Amplification was performed for 45 cycles and the fluorogenic signal was measured during annealing/extension step. Primer set and detecting probe for qRT-PCR was derived from the NP gene of SFTSV: NP forward (5’-CCTTCAGGTCATGACAGCTGG-3’), NP reverse (5’-ACCAGGCTCTCAATCACTCCTGT-3’) and detecting probe (5’-6FAM-AGCACATGTCCAAGTGGGAAGGCTCTG-BHQ1-3’). Copy numbers were calculated as a ratio with respect to the standard control.

### Hematology

To collect hematological data, animals were euthanized and bled by cardiac puncture. Blood was prepared in tubes coated with 0.5 M EDTA (Enzynomics, Daejeon, Republic of Korea). Prior to evaluation of platelet counts, 4% paraformaldehyde was added to EDTA-anticoagulated whole blood samples at a 1:1 ratio to inactivate virus [23]. The platelet counts were analyzed using ADVIA 2012i Hematology System (Siemens Healthimeers, Erlangen, Germany).

### Statistical analysis

Data was analyzed using the Graph Pad Prism 5.01 software (GraphPad Software, La Jolla, CA, USA). Statistical analysis was performed using two-tailed Student’s *t*-test with 95% confidence interval or one-way analysis of variance (ANOVA) followed by Newman-Keuls *t*-test for comparisons of values among different groups. Data are expressed as the mean ± standard deviation (S.D.). Statistical analysis on survival rates were performed using the Mantel-Cox Log Rank test. A *p*-value of < 0.05 was considered statistically significant.

## Results

### Characterization of SFTS DNA vaccines and their gene expression

Four viral genes of SFTSV were cloned into pGX27 vector to be expressed in mammalian cells (Fig. 1). Extracellular domains of viral glycoproteins, Gn and Gc, were separately cloned into the plasmid vector under control of CMV promoter and IRES sequence. Full lengths of viral NP and NS genes fused with a linker peptide (GSGSGSGSGSGRA) were also cloned into the vector under control of RSV promoter. All the proteins were fused with the extracellular domain of Flt3L and the signal sequence of tissue plasminogen activator (tPA) in their N-terminus to promote antigen presentation and trafficking of the fusion proteins as previously described [15]. In addition, pSFTSV-IL12 plasmid also encodes murine IL-12α and β to enhance antigen-specific T cell responses [17]. Expression and secretion of the viral antigens and IL-12 in HEK293 cells transfected with the plasmids were confirmed by ELISA and immunoblot analysis (Fig. 2). All the viral antigens were detected in cell culture supernatants, as well as in cellular lysates (Fig. 2B).

**Fig 2.**
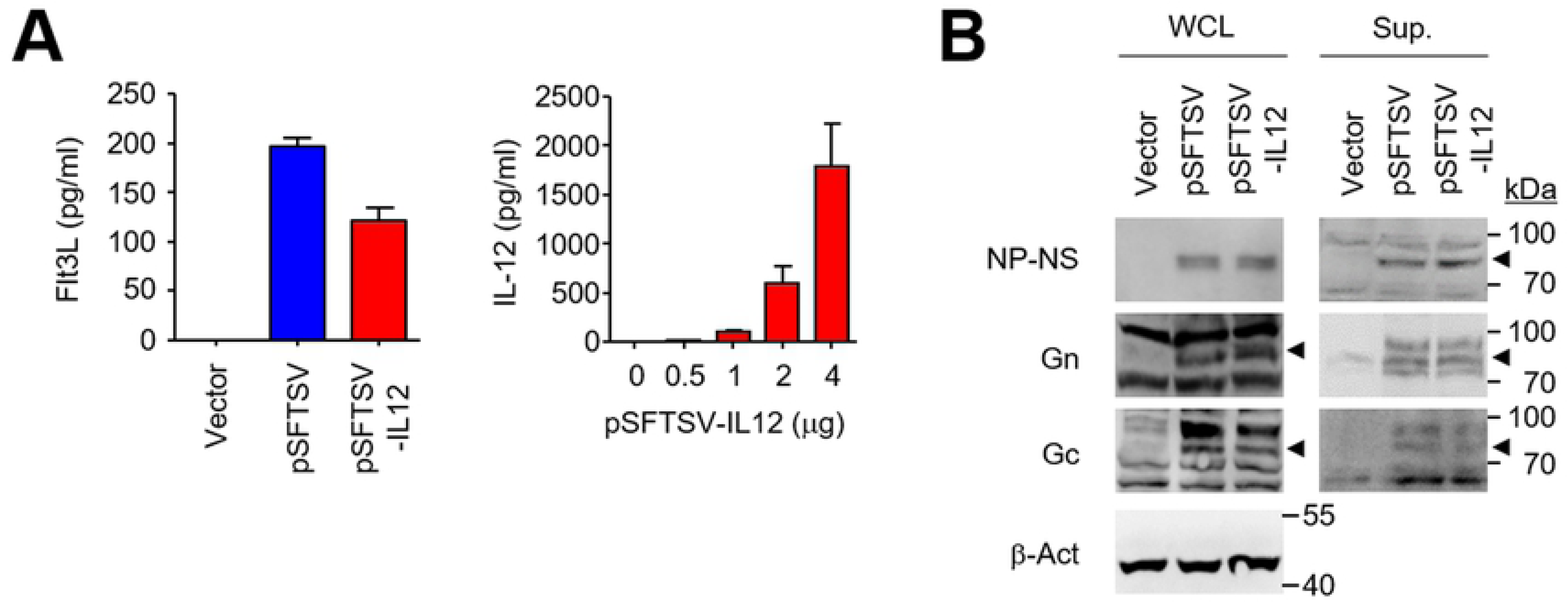
Characterization of gene expression after SFTSV DNA vaccine transfection. (A) Expression and secretion of the viral antigens and IL-12 in HEK293 cells transfected with the DNAs was confirmed by measuring the concentration of Flt3L and IL-12 in the cell culture supernatants by ELISA. Data are presented as mean + S.D. from triplicated experiments. (B) Expression of the viral antigens, Gn (∼ 74 kDa), Gc (∼ 71 kDa), and NP-NS fusion (∼ 80 kDa) proteins, in cell lysates (left panels) and culture supernatants (right panels) of transfected HEK293 cells was assessed by immunoblot analysis using anti-Gn, Gc, or NP antibodies, respectively. The specific bands of antigens corresponding to their expected sizes are indicated with arrow heads. β-actin was used as loading control.

### Antibody and T cell responses against the viral antigens in IFNAR KO mice immunized with the DNA vaccines

Next, we assessed antigen-specific adaptive immunity against the viral antigens after vaccination in IFNAR KO mice. Antibody responses to NP antigen was significantly elevated (mean titer ± S.D.: 255 ± 63, *n* = 3) only in mice immunized with pSFTSV-IL12, and not in mice vaccinated with mock vector or pSFTSV at two weeks after third immunization (Fig. 3A). Specific antibodies against Gn and Gc were barely detectable in all the vaccinated groups (data not shown), suggesting that these DNA vaccines may not efficiently induce specific antibodies against Gn or Gc, but produce NP antibody responses in the presence of IL-12 expression.

**Fig 3.**
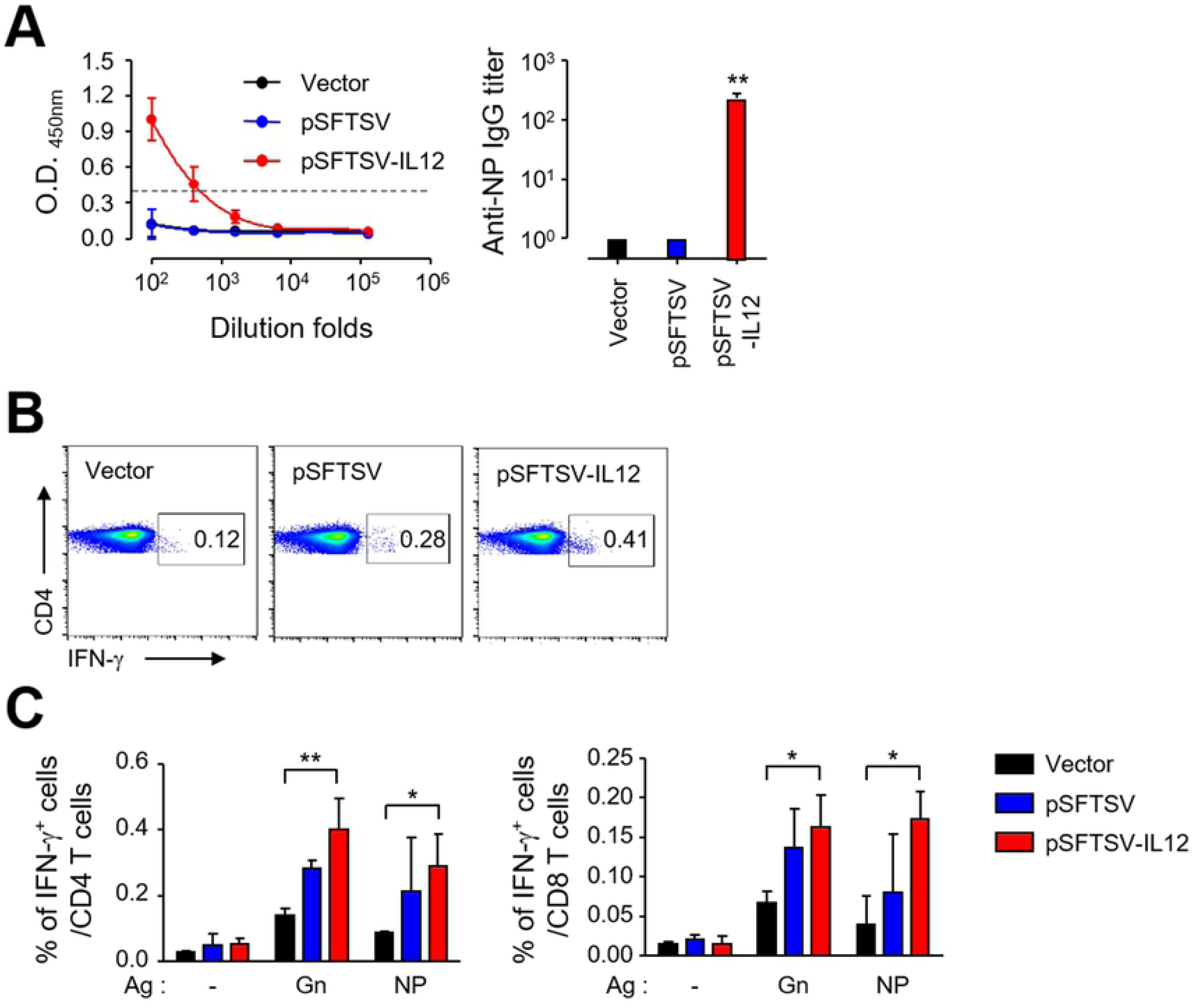
Generation of antigen-specific antibodies and T cell responses in IFNAR KO mice vaccinated with SFTSV DNA vaccines. (A) Anti-NP IgG response was measured by ELISA at two weeks after the third vaccination. The antibody titers of anti-NP IgG in sera of mice (*n* = 3) immunized with different DNA vaccines are presented. Cut-off titers (dashed line, mean O.D. + 3 x S.D. at 1:100 diluents) was determined using sera from vector-immunized mice. Error bar: mean ± S.D. (B and C) Splenocytes were collected from mice at two weeks after the third immunization with the indicated DNA vaccines. Production of IFN-γ by CD4^+^ T or CD8^+^ T cells were analyzed by flow cytometry after stimulation with the indicated antigens. Representative flow cytometric results are presented (B) and the percentile of cytokine positive cells among CD4^+^ or CD8^+^ T cell subsets are summarized (C). Data shown as mean + S.D. from duplicate assays with three mice per group. *, *p* < 0.05; * *, *p* < 0.01.

In order to characterize the potential difference in quality of cell-mediated immunity in IFNAR KO mice immunized with the DNA vaccines, we analyzed antigen-specific T cell responses by assessing IFN-γ secreting T cells in an antigen-dependent manner (Fig. 3B and C). Splenocytes collected from immunized mice were stimulated with Gn or NP antigens and cytokine-positive T cell subsets were analyzed by flow cytometry. The frequencies of cytokine-positive CD4 and CD8 T cells induced by splenocytes of pSFTSV-IL12-immunized mice were significantly higher than those of vector-immunized mice. Even though the cellular responses of immune splenocytes from pSFTSV-vaccinated mice were generally increased, the levels were not statistically significant, suggesting a potent role of IL-12 co-expression in enhancing cell-mediated immunity against the viral antigens.

### Vaccination with pSFTSV-IL12 provides complete protection against lethal infection of SFTSV

Since we observed significant elevation of T cell responses specific to the viral antigens in IFNAR KO mice immunized with pSFTSV-IL12 DNA vaccine, we tested whether it could provide protective immunity against lethal SFTSV infection. The susceptibility of IFNAR KO mice to a Korean SFTSV isolate was examined by inoculating the mice with 1 x 10^1^ to 1 x 10^5^ FFU (S3 Fig A). Upon infection, all the mice gradually lost body weight and became moribund from the third to fifth day after infection, depending on the infection dose. All the infected mice died at 5 ∼ 9 d after infection (S3 Fig A). Similar survival kinetics were previously reported in IFNAR KO mice infected with other SFTSV strains [11, 14], indicating that our Korean SFTSV isolate possesses an equivalent degree of virulence to prior Chinese isolates. The platelet counts in mice infected with 1 x 10^5^ FFU of SFTSV significantly declined up to 4 d after infection, and the mean platelet volumes of the infected mice were significantly higher than those of control animals (S3 Fig B), suggesting that platelet destruction and the activation of platelet production simultaneously occur during lethal infection [14].

To assess the protective efficacy of the DNA vaccine, groups of mice were immunized with mock vector, pSFTSV, or pSFTSV-IL12 three times and then subcutaneously challenged with a lethal dose (10^5^ FFU/mouse) of SFTSV (Fig. 4). All the mock-immunized mice expired by 5 d after SFTSV infection. In contrast, all the mice immunized with pSFTSV-IL12 were protected from lethal viral challenge, and 40% (2 of 5) of mice vaccinated with pSFTSV survived (Fig. 4A). Consistently, pSFTSV-IL12-vaccinated animals lost weight until 4 d after the viral challenge and recovered thereafter, whereas decrease in body weight of pSFTSV-immunized mice was observed up to 8 d after the challenge and the surviving mice gradually recovered. When we examined the progression of thrombocytopenia during the acute phase of infection in the mice, platelet counts in vector-immunized mice rapidly declined until expiration (Fig. 4B, left panel). However, platelet counts only marginally decreased in mice vaccinated with pSFTSV or pSFTSV-IL12 and counts rebounded in pSFTSV-IL12-immunized mice at 4 d after viral infection. In addition, viral loads were significantly reduced in the plasma of pSFTSV or pSFTSV-IL12-vaccinated mice when compared to those of vector-immunized control at 4 d after infection. Notably, despite similar initial viral loads (∼ 10^6^ copies/ml of plasma) at 2 d after viral challenge among the mice groups, pSFTSV-IL12-vaccinated mice (mean ± S.D. = 4.2 × 10^6^ ± 4.6 × 10^6^) were approximately five and fifty times lower than those of pSFTSV-vaccinated mice (mean ± S.D. = 2.1 × 10^7^ ± 1.7 × 10^7^) and vector-immunized mice (mean ± S.D. = 2.0 × 10^8^ ± 1.9 × 10^8^) at 4 d, respectively.

**Fig 4.**
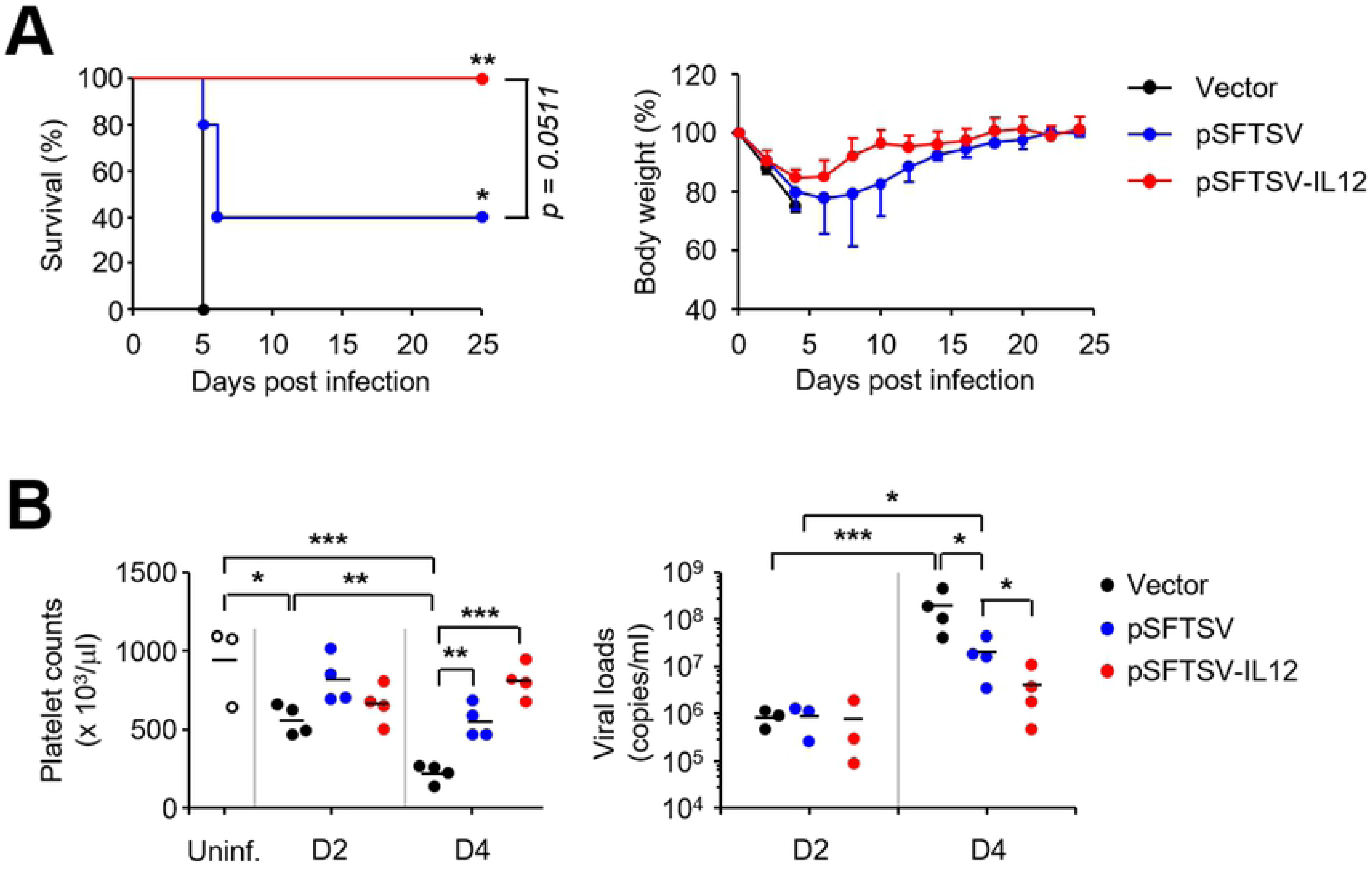
Complete protection of IFNAR KO mice after vaccination with pSFTSV-IL-12. (A) Survival rates (left panel) and body weight changes (right panel) of mice (*n* = 5/group) immunized three times with the indicated DNAs and challenged s.c. with 10^5^ FFU of SFTSV at two weeks after vaccination. (B) Platelet counts (left panel)and viral loads in plasma (right panel) of mice vaccinated with the indicated DNA vaccines are presented. Blood samples were collected at 2 (D2) and 4 (D4) days after viral challenge. Uninf, uninfected; *n* = 3 ∼ 4 mice/group. *, *p* < 0.05; * *, *p* < 0.01; * * *, *p* < 0.001.

### Vaccination of Gn protein provides partial protection against lethal infection of SFTSV

Since antibody responses against the viral glycoproteins, Gn and Gc, were poorly induced in IFNAR KO mice immunized with the DNA vaccines, we further investigate the potential protective role of neutralizing antibodies against Gn or Gc protein by immunization with the protein antigens fused with human Fc fragment (Fig. 5). Vaccination with Gn-Fc or Gc-Fc protein with Alum adjuvant efficiently induced specific antibodies against the immunized antigens in IFNAR KO mice; levels of antibody titers against both antigens were fairly similar (mean titer ± S.D. = 3,584 ± 1,024, *n* = 4). In addition, levels of neutralizing antibodies against SFTSV in immune sera from mice immunized with either protein antigen were significantly higher than those from Fc-immunized controls, when assessed by focus reduction neutralization test (FRNT, Fig. 5B). FRNT_50_ titers of immune sera from mice immunized with Gc-Fc (mean titer ± S.D. = 209 ± 140, *n* = 4) and Gn-Fc (mean titer ± S.D. = 929 ± 1,134, *n* = 4) were approximately five and twenty folds higher than those of Fc-immunized mice (mean titer ± S.D. = 42 ± 29, *n* = 4). It is also notable that FRNT_50_ titers of Gn-Fc immune sera were generally higher than those of Gc-Fc sera, although the difference was not statistically significant. Finally, we examined the protective efficacy of the protein vaccines against lethal infection with SFTSV (Fig. 5C). All the control mice immunized with Fc antigen died at five days after infection, whereas 50% of Gn-Fc immunized mice were protected from lethal challenge and regained their lost body weight. Interestingly, weight loss of Gc-Fc vaccinated mice was delayed and observed from 4 d after infection, whereas the control mice lost weight from 2 d after infection. Even though all the mice immunized with Gc-Fc succumbed to death, they survived three days more than control mice and the extension of median survival time was statistically significant (*p* = 0.0046).

**Fig 5.**
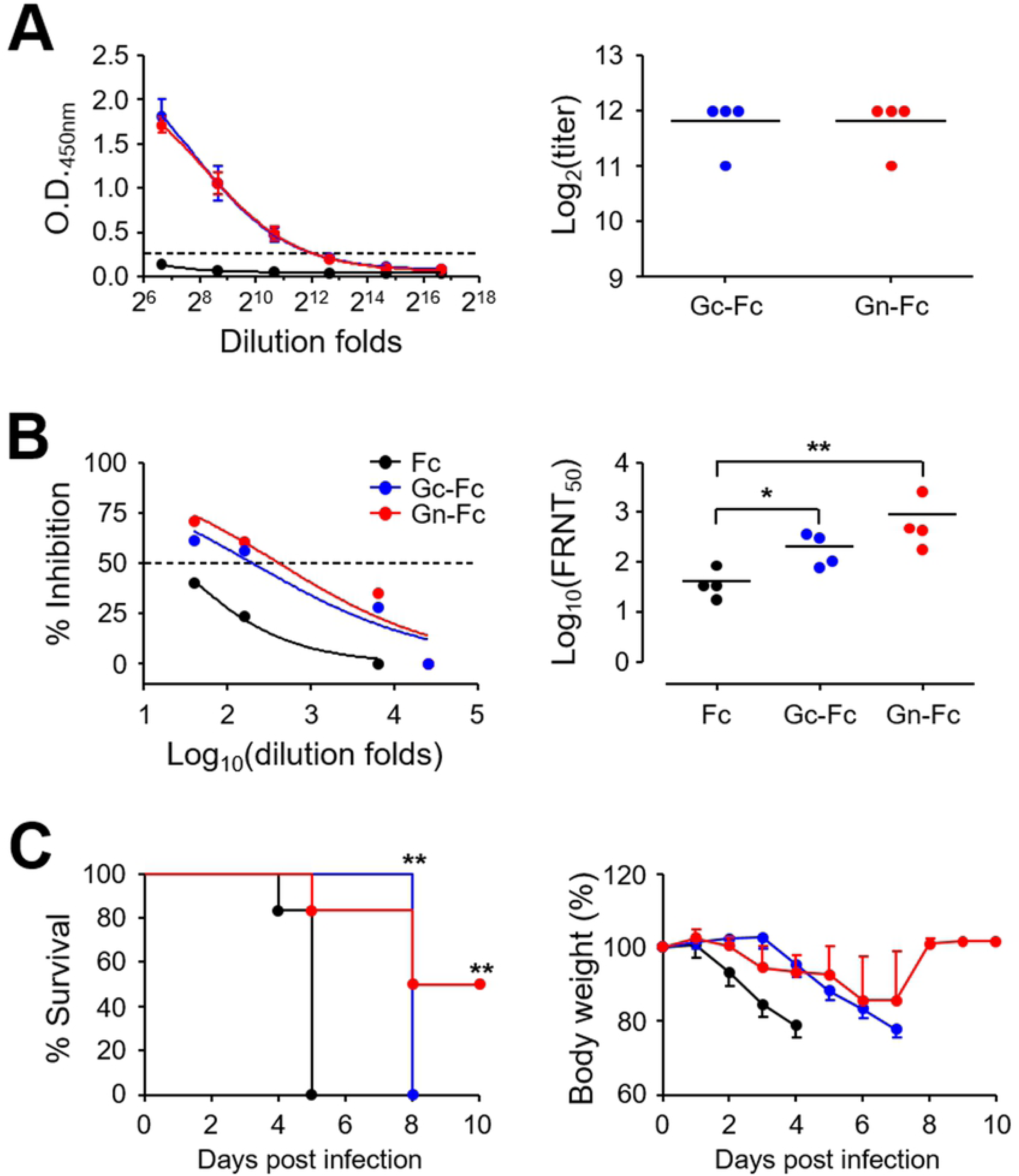
Partial protection of IFNAR KO mice after vaccination with Gn-Fc antigen. (A) Anti-Gc (blue dots) or Gn (red dots) IgG response was measured by ELISA at two weeks after the third immunization with Gn-Fc or Gc-Fc in IFNAR KO mice (*n* = 4/group). Cut-off titers (dashed line, mean O.D. + 3 x S.D. at 1:100 diluents) was determined using Fc-immune sera (black dots). The antibody titers of anti-Gc or Gn IgG in sera of immunized mice are summarized in the right panel. Error bar, mean + S.D.; black line, mean. (B) Neutralizing antibody response to SFTSV generated by SFTSV DNA vaccines in IFNAR KO mice. The amount of neutralizing antibody against SFTSV was determined based on FRNT_50_ (left panel) and summarized (right panel). (C) Survival rates (left panel) and body weight changes (right panel) of mice immunized three times with Fc (black dots, *n* = 6), Gc-Fc (blue dots, *n* = 4), or Gn-Fc (red dots, *n* = 6) antigen and challenged s.c. with 10^5^ FFU of SFTSV at two weeks after vaccination. *, *p* < 0.05; * *, *p* < 0.01.

## Discussion

DNA vaccination has been widely investigated for various infectious diseases, especially targeting viral infections, during the last two decades [24, 25]. Although, human clinical trials of DNA vaccines have yielded poor immunogenicity and less than optimal results, the approval of a few veterinary vaccines is a testimony of proof-of-concept and the hope that licensed DNA vaccines for human use may not be too far away [24]. While we were preparing this manuscript, Kwak J.E. *et al*. reported a promising demonstration of DNA vaccination against lethal SFTSV infection in old-aged (> 4-years-old) ferret model [26]. They used five plasmid DNAs encoding individual viral gene, Gn, Gc, NP, NS, or RdRp, and found that vaccination of old ferrets with a mixture of the five plasmids completely protected ferrets from lethal SFTSV challenge without developing any clinical symptoms and systemic viremia [26]. In addition, adoptive transfer of immune sera from ferrets immunized with two plasmids encoding Gn and Gc transiently induced systemic viremia but provided complete protection against lethal challenge, suggesting that Gn/Gc may be the most effective antigens for inducing protective immunity. They also found that non-envelop (NP, NS, and RdRp)-specific T cell responses also contribute to protection against SFTSV infection although it failed to induce neutralizing activity [26]. Here we also observed that vaccination with a plasmid encoding Gn, Gc, NP, and NS, together with murine IL-12, could provide protective immunity in IFNAR KO mice against lethal SFTSV challenge. However, our current DNA vaccine failed to suppress initial systemic viremia and weight loss (Fig. 4). This might be due to the lack of type I interferon signaling and/or antibody responses against Gn and Gc glycoproteins, which are required for viral neutralization during the early stage of viral infection. Inefficient generation of antibodies against Gn and Gc might be due to the absence of type I interferon signaling in IFNAR KO mice or fusion of Flt3L with the glycoproteins, resulting in conformation defect of the antigens and/or dysregulation of B cell responses. Given that immunization with Gn-Fc and Gc-Fc proteins can induce specific antibodies in IFNAR KO mice (Fig. 5), the absence of type I IFN signaling might not be the cause of inefficient antibody generation. Instead, Flt3L may suppress specific antibody responses as observed in other studies [16, 27-29]. Flt3L may exert tolerogenic effect on CD4 T cells via dendritic cells, thereby promoting B cell hypo-responsiveness *in vivo*; however, the underlying mechanisms of CD4 T cell dysregulation are still unclear [28]. Nevertheless, Flt3L can significantly enhance CD8 T cell responses when fused with a target antigen and promote persistent maintenance of antigen-specific CD8 T cells *in vivo* [16, 29]. In this study, we also observed significant enhancement of Gn and NP-specific CD8 T cell responses, as well as CD4 T cells, in the presence of IL-12 expression (Fig. 3B and C), which correlated well with vaccine protective efficacy in mice infected with lethal doses of SFTSV. The potent effect of antigen-specific T cell responses in protection against SFTSV infection is consistent with the results of the aforementioned ferret infection model [26].

Finally, we evaluated the degree of protective immunity provided by the individual glycoprotein, Gn or Gc, after protein antigen immunization for the first time (Fig. 5), since previous studies showed that vaccination with both antigens retained in a live viral vector [11] or DNA vaccine [26] can provide complete protection against lethal SFTSV challenge in IFNAR KO mice or old ferrets, respectively. Both protein antigens are immunogenic and induce significant levels of neutralizing activity against SFTSV, although anti-Gn antibodies retain relatively higher neutralizing capacity than anti-Gc antibodies. Consistently, the protective efficacy of Gn protein vaccination is relatively higher than Gc protein immunization. While vaccination with either protein antigen failed to provide complete protection against lethal SFTSV challenge, Gc vaccine prolonged the survival time of infected mice and Gn antigen can provide partial protection. Considering that Gn and Gc glycoproteins function as viral ligands for cellular receptor binding and viral membrane fusion in host endosome, respectively [30-32], neutralization of both antigens might be required to completely block viral attachment and entry into host cells.

Taken together, we confirmed that antigen-specific T cell responses against SFTSV induced by DNA vaccination can provide complete protection against lethal viral challenge and immunization with each individual glycoprotein of SFTSV can also confer partial protective immunity. Given that the Achilles heel of DNA vaccines remains their poor immunogenicity in human trials when compared to protein vaccines [25], optimal combinations of DNA and/or protein vaccines, proper selection of target antigens, and incorporation of efficient vaccine adjuvant, need to be further investigated for development of the best SFTSV vaccine formulation for human application.

## Supporting information

**S1 Fig.** Purified Gn-Fc and Gc-Fc proteins were resolved by SDS-PAGE and stained with Coomassie blue.

**S2 Fig.** Gating strategies for T cell analysis by flow cytometry.

**S3 Fig.** Clinical manifestations in IFNAR KO mice infected with SFTSV. (A) IFNAR KO mice were subcutaneously infected with different doses (0, 10^1^, 10^3^, or 10^5^ FFU/mouse) of SFTSV. The mice were monitored daily to assess survival rate (left) and body weight change (right) up to 10 d after infection when all the infected mice died. (B) The blood from the infected mice were collected and platelet counts and their volumes were examined by hematological analyzer. Red line: mean value, *: *p* < 0.05, * *: *p* < 0.01, compared with mock-infected controls.

### Acknowledgements

JGK, KJ, HC, YK, HIK, and HJR received a scholarship from the BK21-plus education program provided by the National Research Foundation of Korea. The funder had no role in study design, data collection, and analysis, decision to publish, or preparation of the manuscript.

## Author Contributions

Conceptualization: JGK KHL NHC.

Formal analysis: JGK KJ HC YK HIK HJR NHC.

Methodology: JGK KJ HC YK HIK HJR YKJ.

Resources: JC KHL.

Supervision: YSK KHL NHC.

Writing - original draft: JGK KJ NHC.

Writing - review & editing: NHC.

